# Interrogating the function of bicistronic translational control elements to improve consistency of gene expression

**DOI:** 10.1101/2023.02.09.527918

**Authors:** Zachary Jansen, Sophia R. Reilly, Matan Lieber-Kotz, Andrew Z. Li, Qiyao Wei, Devon L. Kulhanek, Andrew R. Gilmour, Ross Thyer

## Abstract

Context independent gene expression is required for genetic circuits to maintain consistent and predicable behavior. Previous efforts to develop context independent translation have leveraged the helicase activity of translating ribosomes via bicistronic design translational control elements (BCDs) located within an efficiently translated leader peptide. We have developed a series of bicistronic translational control elements with strengths that span several orders of magnitude, maintain consistent expression levels across diverse sequence contexts, and are agnostic to common ligation sequences used in modular cloning systems. We have used this series of BCDs to investigate several features of this design, including the spacing of the start and stop codons, the nucleotide identity upstream of the start codon, and factors affecting translation of the leader peptide. To demonstrate the flexibility of this architecture and their value as a generic modular expression control cassette for synthetic biology, we have developed a set of robust BCDs for use in several *Rhodococcus* species.

---

Modular and standardized genetic parts are a cornerstone of synthetic biology, enabling rapid and predictable construction of genetic circuits^1–3^. One of the most ubiquitous are translational control elements, which are essential for any genetic circuit that requires protein production. Shine-Dalgarno sites (SD), an element of ribosome binding sites (RBS), are the most common form of translational control element in prokaryotes but are difficult to implement in a format that is both predictable and modular because of variable mRNA secondary structure. SD sites are situated only a few bases upstream of the start codon, and this proximity leads to highly divergent genetic contexts when the same SD sequence is used to control translation of different genes. Different genetic contexts can lead to variable local mRNA structure and the degree to which this structure may occlude the SD can significantly affect the translation of downstream genes^4,5^. A variety of computational tools, most notably the RBS calculator, have been developed which can accurately predict the expression level of a gene from a given RBS, identify problematic local secondary structure, or generate bespoke series of RBSs of varying strengths to tune expression of a target gene^6^. While these tools have revolutionized the design of modular genetic elements for controlling translation, they lack modularity and require unique translational control elements for each gene or genetic context. When optimizing genetic circuits through rapid prototyping with many different genes and contexts, this requirement can become impractical. Bicistronic design translational control elements (BCDs) have previously been proposed to address this limitation and can maintain consistent levels of translation across varying genetic contexts. However, an incomplete understanding of the design rules for BCDs has slowed their adoption in bacterial modular cloning systems.

The earliest reported development of BCD architecture identified three essential elements; (1) A first SD site (SD1) with a fixed genetic context that ensures efficient translation of a short leader peptide or truncated gene, (2) a second SD site (SD2) located within the leader peptide which modulates translation of a gene of interest (GOI), and (3) the stop codon for the leader peptide and a start codon for the GOI located in close proximity 3’ of SD2 which respectively serve to terminate translation of the leader peptide and initiate translation in the correct reading frame for the GOI (**Figure 1 and S1**)^7^. The initial designs were improved by pairing the empirically validated *trpL* leader peptide and 5’ UTR with a second SD site and translation start site from the *E. coli trpE-trpD* junction^8,9^. Interrogation of these newer designs established that helicase activity of the ribosome played an important role in effective translation of the downstream gene. Later efforts to develop a series of modular translational control elements adopted these scaffolds and generated a series of SD2 variants which spanned a 600-fold range of reporter protein expression^10^.

**Figure 1:**
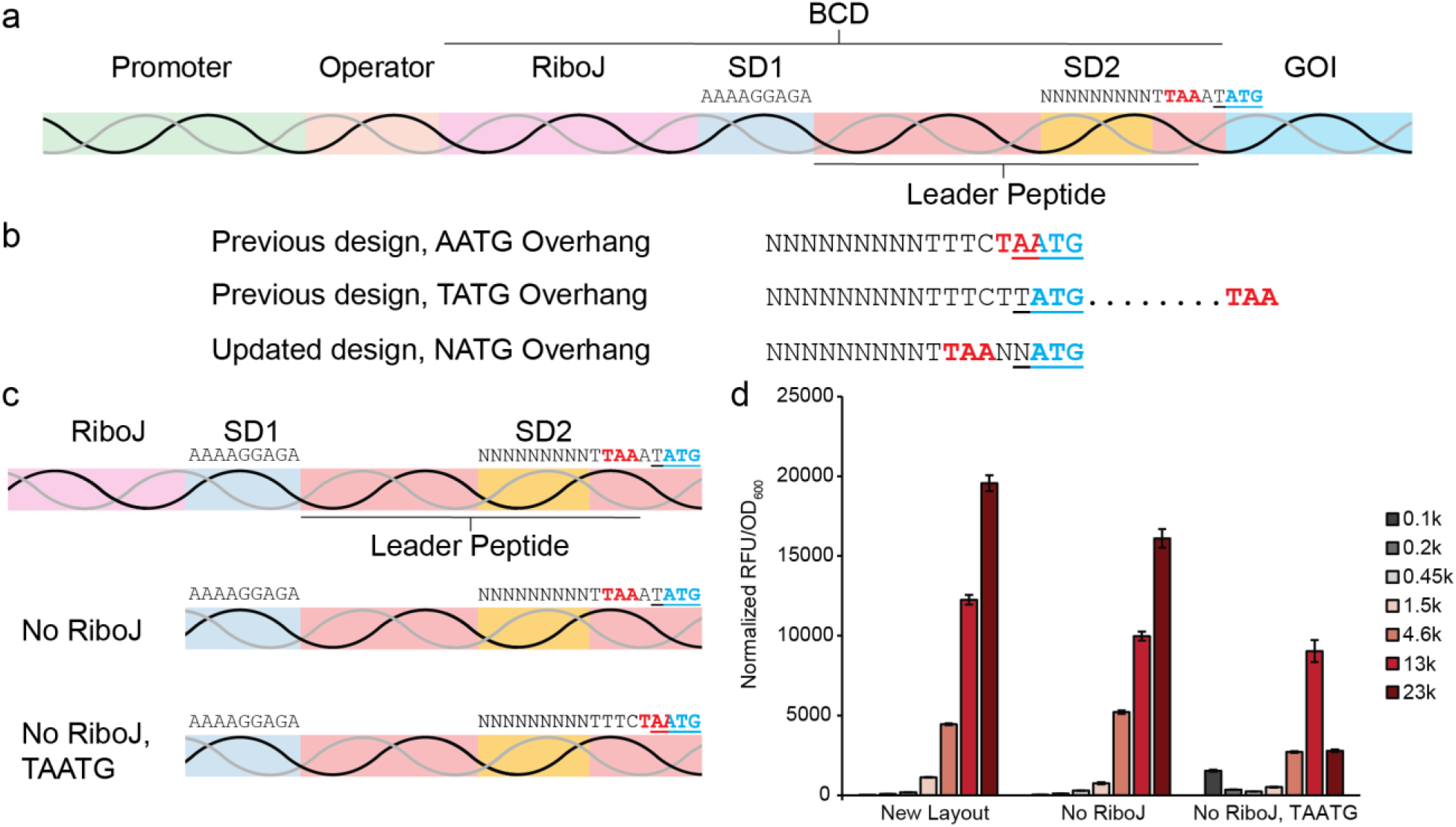
SD sequences are specific for different ribosomal reinitiation motifs. (a) The new layout of the bicistronic translational control element is shown in a modular framework and the DNA sequences of the Shine-Dalgarno sites and the ribosomal reinitiation region surrounding the stop and start codons is indicated. (b) DNA sequences of the previously reported and updated translation reinitiation region are depicted with either an AATG or TATG cloning overhang. The start and stop codons are marked in blue and red respectively and the 4 nt overhangs are underlined. (c) The layout of three different BCD designs, the first including RiboJ and separated stop and start codons in the ribosomal reinitiation region, the second lacking RiboJ, and the third lacking RiboJ and featuring an overlapping stop and start codon. (d) Fluorescence assay using a subset of newly derived SD2 sequences transferred into the three different BCD layouts.

This architecture has been successfully adapted for use in several bacterial species and proven to be a valuable tool for synthetic biology by yielding highly predictable, rank order expression across diverse genetic contexts^11–15^. However, important details regarding the mechanism of translation initiation at SD2 remain unclear. In particular, our incomplete understanding of the relative contributions of ribosomal reinitiation at SD2 versus recruitment of free 30S subunits and sequence elements affecting this process complicate efforts to improve the current BCD architecture beyond further randomization of the SD2 region.

Another limitation is the use of an overlapping TAATG motif in the previously reported BCD series which encodes both the stop codon of the leader peptide and the start codon of the downstream gene. This motif restricts BCD compatibility to modular cloning systems which use the AATG overhang upstream of modular CDS parts. This excludes parts compatible with modular cloning systems using the common TATG overhang or specific use cases where a CATG or GATG overhang is required. Our initial efforts to separate the overlapping stop and start codons in the TAATG motif by moving the TAA codon one position upstream, a seemingly conservative modification which does not affect the relative positions of SD2 and the start codon and retains two T nucleotides immediately downstream of SD2, completely abolished the rank order of previously reported BCDs and raised further questions about the mechanism of this design.

In this work we modify the BCD architecture by adjusting the spacing between the stop and start codons in the ribosomal reinitiation region to allow for diverse toolkit overhangs and incorporating a self-cleaving ribozyme to the 5’ end of the BCD to improve standardization across different transcriptional contexts. We derive a new series of SD2 sequences within this modified BCD architecture that span nearly three orders of magnitude translational strength and enable highly reproducible low-level translation initiation for the first time. We further show that, in addition to *E. coli*, these design changes can function well in both model and non-model gram positive *Rhodococcus* species.

## Results and Discussion

A modified BCD architecture was designed with elements to render it agnostic to different modular cloning systems and to standardize the 5’ end. To separate the stop and start codons and allow for alternate cloning overhangs the stop codon was moved two bases upstream of the start codon, which shortens the leader peptide by one amino acid and results in a sequence of TAANNATG (**Figure 1a-b**). We use the sequence TAAATATG for all future experiments unless otherwise noted. Furthermore, we added the self-cleaving ribozyme RiboJ to the 5’ end of the BCD. Cleavage of the mRNA transcript by RiboJ leads to a consistent 5’ sequence and structure which serves to insulate it from the upstream genetic context, as well as stabilize the mRNA via the 23 nt RiboJ hairpin (**Figure S2**)^16,17^.

A subset of the SD2 sequences from Mutalik *et al*. which span the reported ~600-fold dynamic range (BCDs 2,7,8,13,16,17,18, 22, and 23) were transferred into this new layout^10^. Plasmids were assembled with a medium strength promoter (P150) and a BCD variant controlling expression of the red fluorescent protein mScarlet-I^18^. We observed that these sequences did not maintain the previously reported rank order in the new BCD configuration and displayed only ~10-fold dynamic range (**Figure S3**). Additionally, in this genetic context the highest observed strength BCD (BCD13) resulted in slow growth and lower final cell densities indicative of toxicity. These data suggest that in order to extend the dynamic range, weaker BCDs were required. The previously reported BCD series conserves a central three-nucleotide core in the nine-nucleotide SD2 region which is not randomized and corresponds to the consensus SD sequence. While this undoubtedly facilitated the isolation of functional SD2 sequences, it likely limited the dynamic range of the previously reported BCD series. To overcome the incompatibility of the previous SD2 sequences with the new BCD sequence context, we derive a new set of SD2 sequences that span a broader range of desired translation strengths.

Initially, a library encoding 4^9^ SD2 variants was generated upstream of mScarlet-I using degenerate oligos which randomized the entire nine bp SD2 region. From this library nine BCDs were isolated (0.1k, 0.75k, 2.3k, 4.6k, 8.5k, 13k, 18k, 23k, 25k) spanning over 800-fold dynamic range (**Figures 1d, 2a**). A small pool of BCD variants with strengths exceeding 25k were also isolated, however, in the context of our constitutively expressed reporter we observed significant toxicity. Importantly, the strongest BCD derived in this context, 25k, has a slight impact on growth rate indicating an effective ceiling on translational initiation rate. Experiments to identify stronger SD2 sequences found only a few candidates leading to slightly stronger expression (supplementary results). Preliminary use of the new series of BCDs revealed a need for additional gradations at the low end of the scale, which would also serve to effectively reduce translational bursting^19,20^. We attempted to enrich for weak SD2 sequences by reconstructing the mScarlet-I reporter downstream of a much stronger constitutive promoter (P500) and repeating the SD2 library screen^18^. This approach yielded a large number of moderate strength SD2 sequences, and just a single new weak BCD variant (1.5k). Adopting the approach used by Mutalik *et al*., whereby a partial SD2 core was retained during library screening, we used the weakest isolated SD2 sequence from BCD 0.1k as a scaffold and randomized three 3-nt blocks which were pooled for screening. While no candidates were recovered with weaker expression than 0.1k, two new weak SD2 sequences were identified, 0.45k, and 0.2k. Complete sequences for the *E. coli* BCD series can be found in **SI Table 1**.

**Figure 2:**
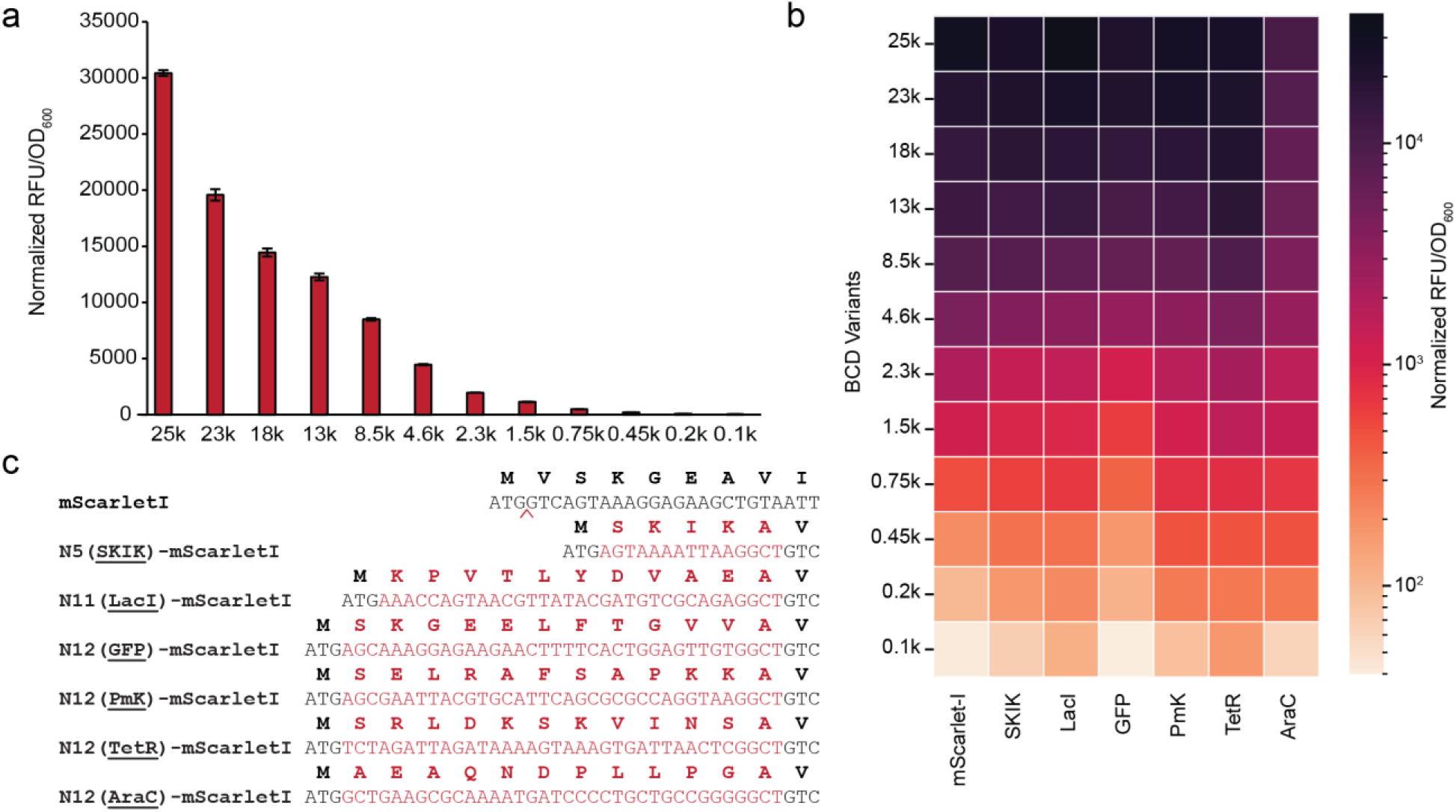
BCDs maintain consistent translational strengths across diverse genetic contexts. (a) Relative expression of mScarlet-I using the complete series of BCD translational control elements. (b) Heatmap showing the relative fluorescence of mScarlet-I using the series of BCDs in seven different genetic contexts around the translation initiation region. (c) The DNA and amino acid sequences of each different genetic context used to evaluate the consistency of reporter expression. Amino acid tags were inserted between the ATG start codon and the second amino acid of the mScarlet-I gene. Fluorescence was normalized between contexts using BCD 8.5K and wild-type mScarlet-I as an internal control which is set to 8500 RFU/OD_600_.

A subset of this BCD series (0.1k, 0.2k, 0.45k, 1.5k, 4.6k, 13k, and 23k) was selected for investigation into which changes to the BCD architecture affected performance of the previously reported SD2 sequences. This reduced BCD series was evaluated in the absence of RiboJ, and when both RiboJ is removed and the original TAATG reinitiation motif was restored (**Figure 1c**). While removal of RiboJ alone had no effect on performance of the BCD series, restoration of the original start and stop codon spacing abolished the previously observed rank order and greatly reduced the dynamic range (**Figure 1d**). This result indicates that while BCDs maintain translation rates across genetic contexts downstream of the start codon, changes to the ribosomal reinitiation region can have a significant effect.

The critical feature of the BCD design is their interoperability between different genetic sequences flanking either side of the start codon^10^. To determine if the newly derived BCD series maintains this functionality, we evaluated their performance across seven different genetic contexts with regard to translation strength and preservation of the rank order (**Figures 2b, S4**). The first 12 amino acids of five different genes were each appended to the N-terminus of mScarlet-I, generating five different genetic contexts. A sixth context appended the five-amino acid SKIKA tag, which has been shown to improve ribosomal procession^21^. In the six contexts with N-terminal tags, the additional nucleotides were added between the start codon and the second codon in mScarlet-I without duplicating ATG codons. The native mScarlet-I N-terminus served as the seventh genetic context. The nucleotide and amino acid sequences of these seven genetic contexts are depicted in **Figure 2c**. We observed that the rank order of the 12 BCD variants is maintained across all genetic contexts tested, with minor variation in the relative fluorescence.

Our initial changes to the BCD design were prompted by the need to construct BCDs that are compatible with multiple different modular golden gate overhangs. To determine whether our new design maintains equivalent transcriptional rates across variable modular cloning overhangs we mutated the modular overhang from the wildtype TATG to AATG, CATG, and GATG. Additionally, we mutated the two nucleotides flanking the stop codon to determine the effect of changing the context around the stop and start codons. These mutations had a minor effect on fluorescent protein output (**Figure S5**). To investigate the effect of differential spacing between the stop and start codon, we constructed four variants which fixed the relative positions of the start codon and SD2 but encoded leader peptides of different lengths (**Figure S6**). To accommodate a stop codon one position further upstream than our design, the final two nucleotides of the SD2 sequence were fixed as TG. Using the 0.75k SD2 sequence as a scaffold, this new variant was empirically validated and designated 3.75k. In contrast with our previous observations, only slight differences in mScarlet-I fluorescence were observed between the different spacings using this SD2 sequence. Surprisingly, this included the overlapping ATGA motif which has been shown to result in strong, SD independent ribosomal reinitiation^22^. Taken together, these conflicting results suggests a complex, and context dependent relationship between SD2 strength and spacing of the start and stop codons.

It is well established that in a bicistronic layout, either with full-length genes in native contexts or artificial leader peptides, complete translation of the upstream gene is essential for translation of the downstream gene^23^. However, the relative contributions of ribosomal initiation at SD1 followed by reinitiation at SD2 versus free ribosomal subunits assembling and initiating translation at SD2 are unclear. To investigate this phenomenon, we built several new BCD designs (**Figure 3**). We first eliminated the stop codon of the leader peptide from the ribosomal reinitiation region. This resulted in ribosomes that initiated translation at SD1 continuing into the coding sequence of mScarlet-I thereby adding an additional nine amino acids to the leader peptide before reaching the next in-frame stop codon. This design resulted in greatly decreased fluorescence, even though the SD2 sequence was unaltered and should lack secondary structure due to the passage of ribosomes translating the leader peptide. This indicates that the vast majority of ribosomes begin translation at SD1 and reinitiation is the primary mechanism behind translation in the downstream gene. One explanation for this observation may be that ribosomes which initiate translation at SD1 sterically obstruct ribosomal assembly at SD2. To test whether this is the case, we constructed a BCD variant where the SD1 site is mutated to CCCCCCCCC. This poly-C sequence in place of the original SD1 sequence was confirmed to poorly initiate translation when empirically validated in a mono-cistronic design. When SD1 is mutated to poly-C in the BCD, we observe no increase in fluorescence as compared to the original SD1 sequence. This indicates that ribosomes initiating translation at SD1 are not blocking ribosomal assembly at SD2, and that termination of leader peptide synthesis sufficiently far downstream of the reinitiation region prohibits translation of the second gene. This observation was consistent across several different promoter strengths, although stronger promoters were incompatible with the native SD1 sequence in a mono-cistronic layout.

**Figure 3:**
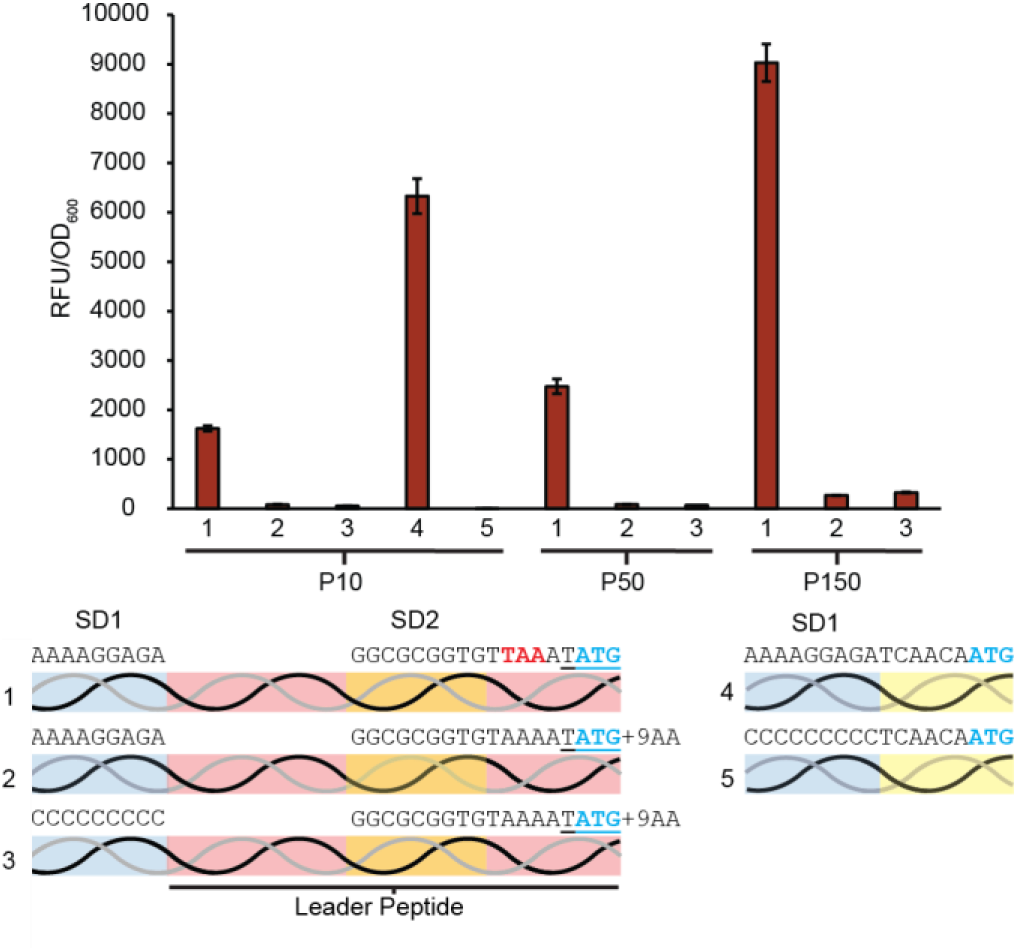
Free ribosomes play a minimal role in the translation of genes downstream of BCDs. Eliminating the stop codon in the leader peptide (2) enables the ribosome to progress beyond the SD2 region and start codon before terminating, resulting in minimal translation of the downstream gene. Eliminating translation from SD1 by using a poly-C sequence (3), thus enabling free ribosomal subunits to assemble at SD2 without ribosomes translating the leader peptide blocking access to SD2 does not restore gene expression. This relationship was consistent across three different promoter strengths. Translational strength of the different SD1 regions was confirmed in a monocistronic design (4 and 5).

Following their development and utilization in *E. coli* there have been several reports of successful transfer of BCDs into other bacteria^12,15^. To confirm that the new BCD architecture developed here is equally portable, we developed a small series of BCD variants for *Rhodococcus*, an industrially relevant genus of gram positive Actinomycetota^24^. To construct these BCDs, we first replaced the SD1 sequence with a previously reported SD sequence (ACTAAGGA) found in *Rhodococcus* that resulted in high levels of translation^25^. The leader peptide sequence was changed to the first 12 amino acids of the *E. coli lacZ* gene, which has been empirically validated with the ACTAAGGA SD1 and shown to yield strong expression^25^. This design was evaluated in combination with the second strongest SD2 sequence from the *E. coli* series (23k) and the far-red fluorescent protein E2-Crimson in *Rhodococcus ruber* C208, a non-model environmental *Rhodococcus* species. The 23k SD2 sequence yielded moderate fluorescence and we subsequently randomized the entire SD2 region to identify an additional range of BCD strengths. Following screening in *R. ruber* C208 we isolated a small pool of BCDs spanning a >250-fold dynamic range. This series maintains the expected rank order of translational strengths in two additional *Rhodococcus* species, *R. erythropolis* N9T-4 (NBRC 110906) and *R. imtechensis* (DSM 45091) (**Figure 4 and S7**). This data illustrates that our new BCD architecture can function robustly across species when using an empirically validated SD1-leader peptide pairing.

**Figure 4:**
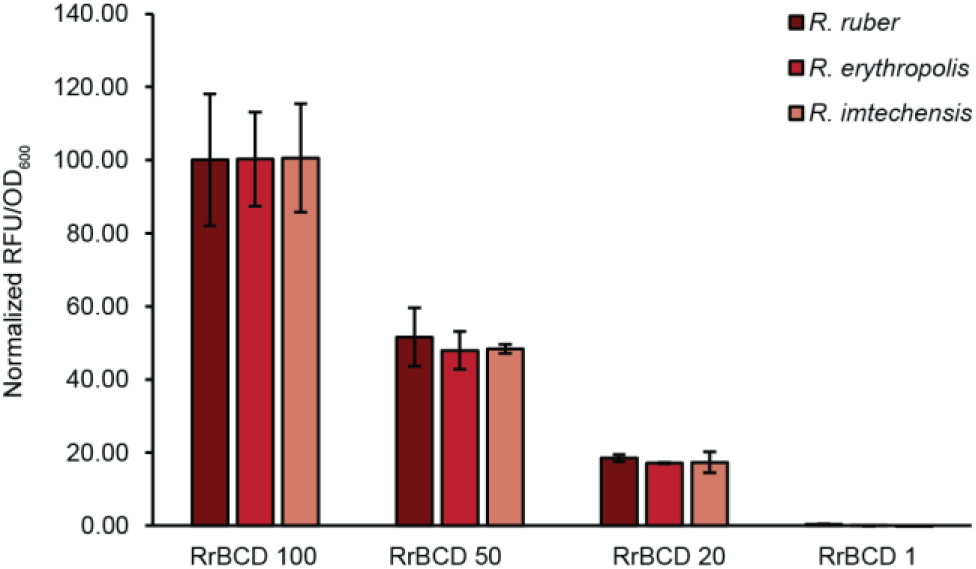
BCDs retain their relative rank order in different *Rhodococcus* species. Relative fluorescence of E2-Crimson regulated by four *Rhodococcus* BCD variants in three different species. Data is normalized between species such that RrBCD100 in each species is set to 100 RFU’s, and each subsequent BCD is normalized to this value. Raw values vary across species and are shown in **Figure S7** without normalization.

In this work we have developed a new BCD architecture that is compatible with modular cloning frameworks using a variety of common cloning overhangs incorporating an ATG start codon. We show that the SD2 sequences are not transferable between translation reinitiation contexts and that the newly derived SD2 sequences preserve the most important aspect of the BCD design, the stability of translation initiation rate across genetic contexts. We have further investigated which nucleotide region contributes to these effects and demonstrate that BCDs function almost exclusively via translation reinitiation rather than free ribosome assembly at SD2. We anticipate these results will prove useful to researchers needing robust translational control elements for high-throughput prototyping of genetic circuits and those developing BCDs for additional non-model bacteria in the future. Based on this work, we also identify several potential avenues to further improve the BCD architecture including, (i) leader peptide designs which impose a lower metabolic burden via different amino acid usage or more efficient recycling, (ii) identifying the minimum length for an effective leader peptide, (iii) the introduction of amino acids motifs at the N-terminus of the leader peptide which facilitate efficient transition of the ribosome into the elongation phase through reducing stalling or abortive cycling, and (iv) development of BCDs with improved sequence diversity

## Materials and Methods

### Bacterial strains and growth conditions

*Rhodococcus ruber* C208 (DSM 45332), *Rhodococcus erythropolis* N9T-4 (NBRC 110906) and *Rhodococcus imtechensis* (DSM 45091) (syn. *R. opacus)^26^* were cultured at 28 °C in Tryptic Soy Broth (TSB) supplemented with kanamycin at a concentration of 100 μg/mL for *R. ruber* and *R. imtechensis*, and 200 μg/mL for *R. erythropolis. E. coli* strain DH10B was used for plasmid construction and all experiments other than those evaluating BCD performance in *Rhodococcus* species. *E. coli* strain DH10B was cultured at 37 °C in Terrific Broth supplemented with chloramphenicol at a concentration of 33 μg/mL for plasmid maintenance.

### Plasmid construction

All plasmids were constructed using a hierarchical, golden gate assembly system. Assembled plasmids used in *E. coli* are comprised of a pMB1 origin of replication, a chloramphenicol resistance marker, and an expression cassette for mScarlet-I containing a constitutive promoter, the variable translational control elements, the mScarlet-I CDS, and a transcriptional terminator. Plasmids used in *Rhodococcus* sp. are assembled as shuttle vectors containing both a pUC origin of replication and a pNC903 *Rhodococcus* origin of replication, the **aph(3’)-IIa***a* Kanamycin resistance marker, and an E2-Crimson expression cassette consisting of a constitutive promoter, bicistronic translational control element, the E2-Crimson CDS, and a transcriptional terminator.

### Randomized BCD library construction

Construction of the libraries of bicistronic translational control elements was performed by PCR using oligonucleotide primers containing degenerative bases. PCR products were circularized using Gibson Assembly. In *E. coli*, this library was immediately transformed into chemically competent DH10B cells. In *Rhodococcus*, assembled libraries were desalted and then transformed into *Rhodococcus* species via electroporation.

### Electrocompetent cell preparation

Electrocompetent *Rhodococcus* cells were prepared by culturing the bacteria in 200 mL TSB containing 1.5% w/v glycine and 1.5% w/v sucrose. Cells were cultured at 28 °C for 16 h and then washed three times with 20 mL sterile 10% glycerol, split into 200 μL aliquots and flash frozen in liquid nitrogen. Cell aliquots were stored at −80 °C until use.

### *Rhodococcus* electroporation protocol

100 μL aliquots of electrocompetent *Rhodococcus* cells were mixed with 100 ng of plasmid DNA, transferred to 0.2 mm cuvettes, and electroporated at 2.5 kV with a capacitance of 25 μF. Cells were recovered in 2 mL TSB at 28 °C for three hours with 250 rpm agitation, harvested by centrifugation and resuspended in 150 μL TSB, and transferred to petri plates containing solid tryptic soy agar medium supplemented with kanamycin. Plates were incubated at 28 °C for several days until colonies were observed.

### Fluorescence assays

Transformants were initially cultured overnight in terrific broth supplemented with chloramphenicol. The following day, 30 μL of stationary phase culture was added to 970 μL of fresh medium supplemented with chloramphenicol in a 96-well deep-well plate. Cultures were incubated for six hours and then harvested by centrifugation at 3500 x *g* for 10 minutes. Cell pellets were resuspended in 1 mL PBS pH 7.4 and 100 μL of cell suspension was transferred to a 96-well microtiter plate. Fluorescence assays were conducted using a multimode plate reader (Tecan M200 Pro). Absorbance was measured at 600 nm and fluorescence measured using an λ_ex_ 550 nm and λ_em_ 610 nm.

For *Rhodococcus* species, transformants were selected and cultured in 24-well deep-well plates in tryptic soy broth supplemented with kanamycin at 28 °C. Cultures were incubated for three days with constant agitation until reaching stationary phase. Subcultures were started using 50 μL of confluent culture to inoculate 2 ml of fresh medium in 24-well deep-well plates. These cultures were incubated at 28 °C for two days. 500 μL aliquots from each well were centrifuged, and the cell pellets were resuspended in an equal volume of PBS pH 7.4. 100 μL of cell suspension was transferred to a 96 well microtiter plate and assayed by measuring absorbance at 600 nm and fluorescence using an λ_ex_ 595 nm and λ_em_ 650 nm.

### Statistical analysis and data visualization

Unless otherwise indicated, all data was collected from a minimum of three biological replicates using three technical replicates for *E. coli*, and two technical replicates for *Rhodococcus* species. Error bars represent the standard deviation of the biological replicates.

## Data availability

Plasmids encoding BCDs as modular parts for *E. coli* and *Rhodococcus* are available on Addgene.

