## Supplementary Information for "Interrogating the function of bicistronic translational control elements to improve consistency of gene expression"

This file contains:
Supplementary Results
Supplementary Figures 1-8
Supplementary Tables 1 and 2

### Supplementary results

Following our efforts to generate weak BCDs, which indicated that BCD strengths are constrained by the circuit conditions under which they are derived, we sought to further demonstrate this phenomenon in a different circuit context. We transferred our strongest BCD, 25k, into a new context by switching the promoter from a medium strength promoter (P150) to a weak promoter (P10). A new $4^9$ library of SD2 sequences was generated from this template and 352 colonies were randomly selected and screened for fluorescence after 16 hours of growth. We observed that our selected colonies had a dynamic range of fluorescence exceeding 1500-fold (**Figure S8**). This included a small number of SD2 variants which yielded greater fluorescence that BCD 25k. Given BCD 25k demonstrates toxicity when paired with mScarlet-I and promoter P150, it is unlikely these variants would be isolated from a library screened in the context of a stronger promoter. To generate the widest possible range of translational strengths, it is likely necessary to use a series of promoters to capture SD2 diversity at each end of the range.

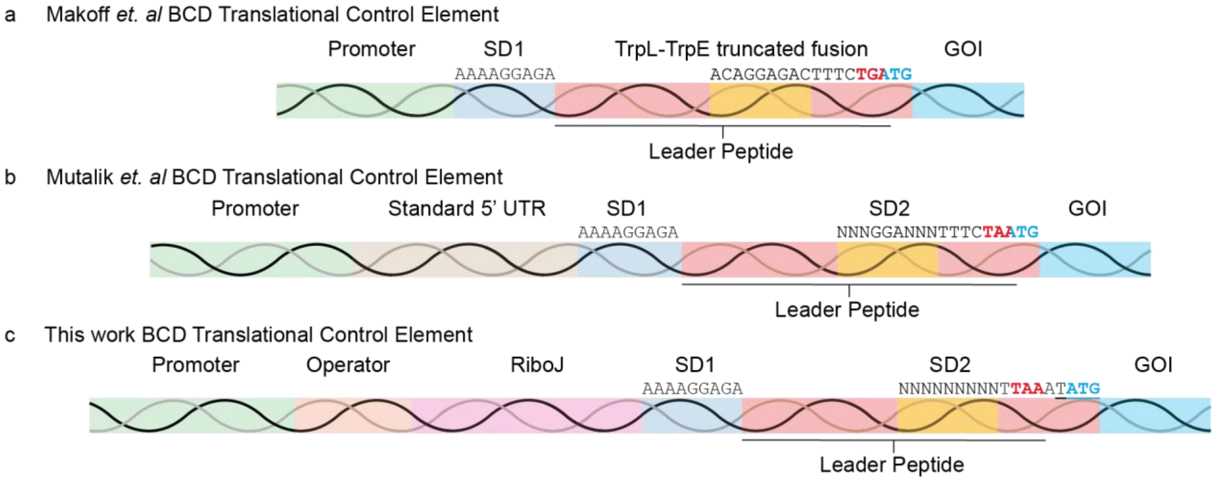


**Figure S1:** Progression of different bicistronic design translation control elements. (a) The first iteration developed by Makoff *et al.* fused the 5' UTR and N-terminus of the *E. coli trpL* gene to the C-terminus from the *E. coli trpE* gene to express a reporter gene. (b) The second iteration developed by Mutalik *et al.* as a synthetic BCD and included a series of SD2 sequences with a central 3 nt GGA motif. The promoter and gene of interest could be changed between constructs. (c) In the third design (this work), the BCDs include a RiboJ self-cleaving ribozyme at the 5´ end to improve the stability and standardization as well as an altered ribosomal reinitiation region which preserves compatibility with a NATG cloning overhang.

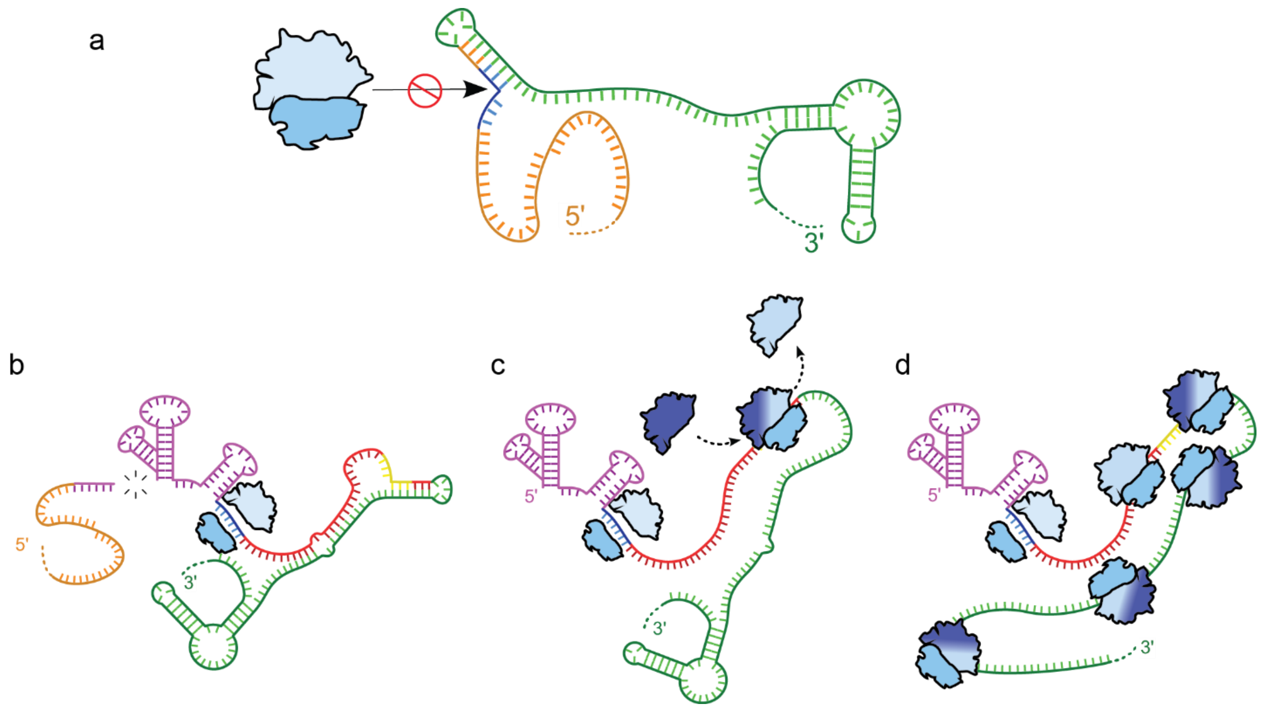


**Figure S2:** Schematic of BCD function. RNA bases are color coded to indicate the different genetic element; 5' untranslated region (orange), RiboJ (purple), SD1 sequence (blue), leader peptide (red), SD2 site (yellow), and the downstream GOI (green). The large and small subunits of ribosomes are colored in blue. (a) A mono-cistronic design ribosome binding site where the SD sequence is occluded by RNA structure. In a mono-cistronic design the RNA can form divergent structures around the SD site in different genetic contexts leading to inconstant availability of the SD sequence for ribosome recruitment. (b) In the new bicistronic design, ribosomes assemble at the first SD sequence which is set in a consistent genetic context, and the RiboJ undergoes self-cleavage leading to a consistent 5' structure and sequence. (c) The helicase activity of the ribosome unwinds the structure of the RNA as it translates the leader peptide. Once reaching the stop codon of the leader peptide the ribosome may terminate translation and release the mRNA, or the small subunit of the ribosome may recognize the nearby start codon for the GOI and the SD2 sequence and reinitiate translation. The original large subunit of the ribosome may be retained, or a new large subunit may bind to the small subunit. (d) Translation of the GOI follows ribosomal reinitiation at SD2 leading to protein production.

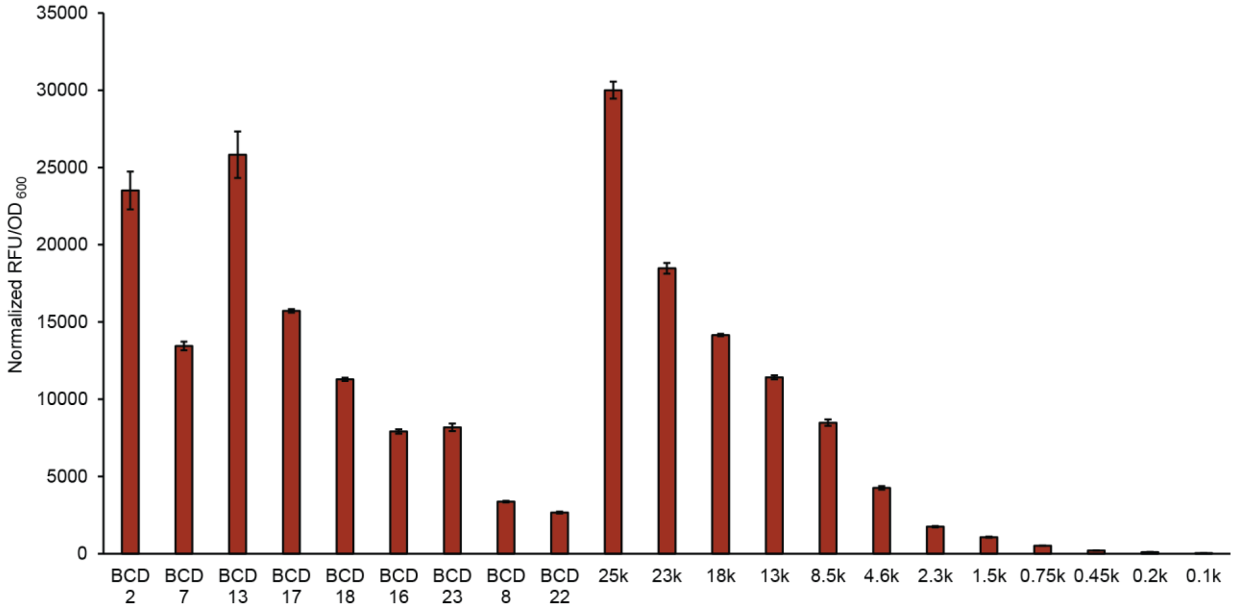


**Figure S3:** SD sequences are not transferable between different BCD layouts. Nine SD2 sequences spanning the complete range of translational strengths previously reported by Mutalik *et al.* were cloned into the new, non-overlapping BCD layout. These SD2 variants are shown in order of reported translation initiation strength. The newly isolated series of BCDs are shown for context. The construct expressing wild-type mScarlet-I using BCD 8.5K was used as an internal reference and data is normalized such that BCD 8.5k expressing mScarlet-I is equivalent to 8500 RFUs. Toxicity, indicated by reduced cell density, was observed for BCD 13 and BCD 25k.

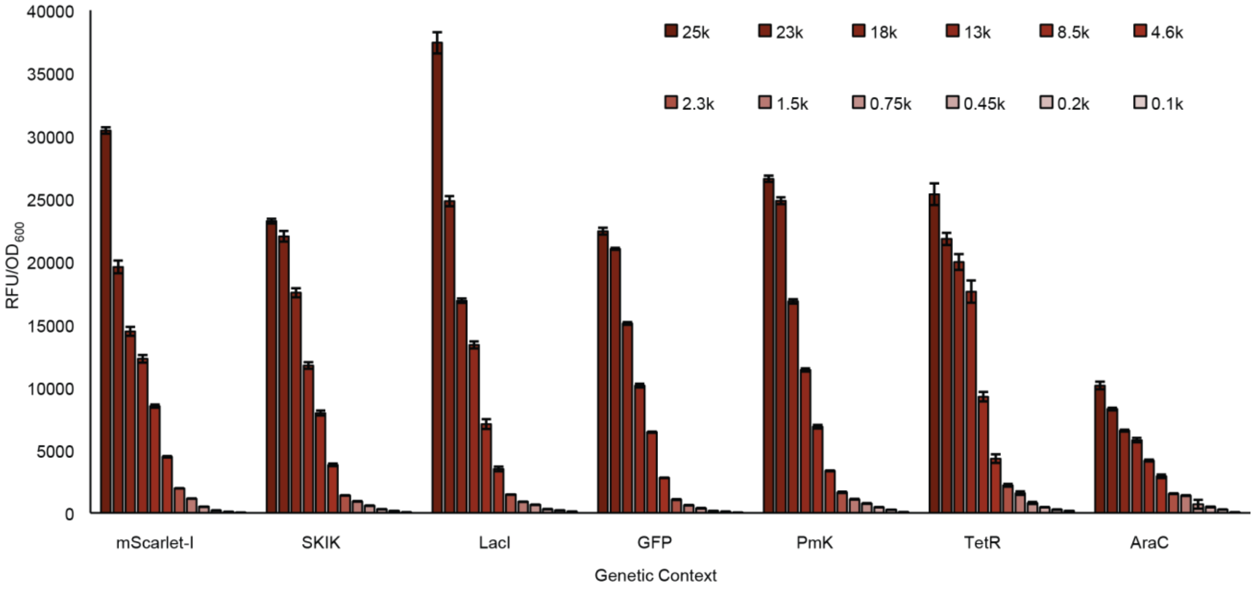


**Figure S4:** Relative fluorescence of each BCD expressing mScarlet-I in seven different genetic contexts. All twelve BCD variants maintain the expected rank order in all seven genetic contexts. The construct expressing wild-type mScarlet-I using BCD 8.5K was used as an internal reference and data is normalized such that BCD 8.5k expressing mScarlet-I is equivalent to 8500 RFUs.

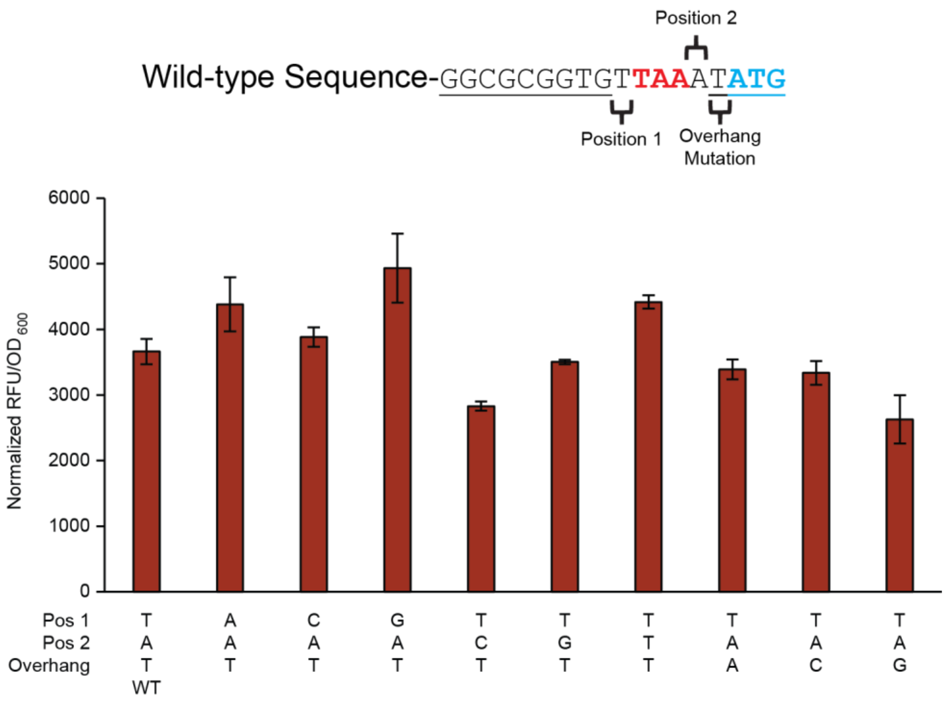


**Figure S5:** Mutations to bases surrounding the ribosomal reinitiation region have variable effects in the context of BCD 8.5k. Wildtype sequence of the ribosomal reinitiation site is shown with positions indicated that were mutated to the other possible nucleotides. The SD2 sequence is underlined.

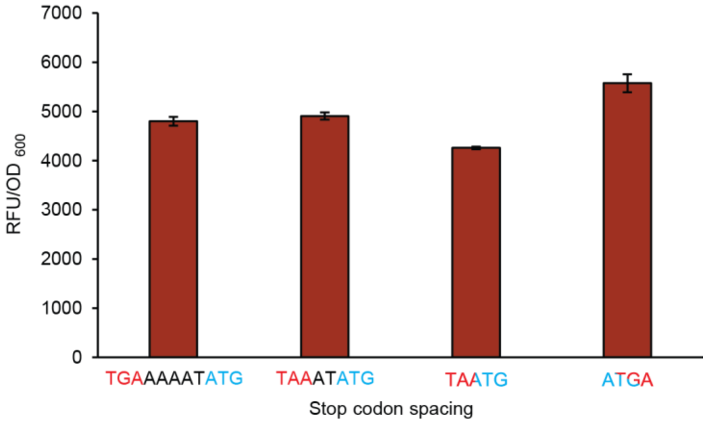


**Figure S6:** Effect of spacing between the leader peptide stop codon and the start codon of the downstream gene with BCD 3.75K. Labels indicate the sequence spanning the stop and start codons. Stop codons are indicated in red, start codons are indicated in blue, and intervening nucleotides are indicated in black.

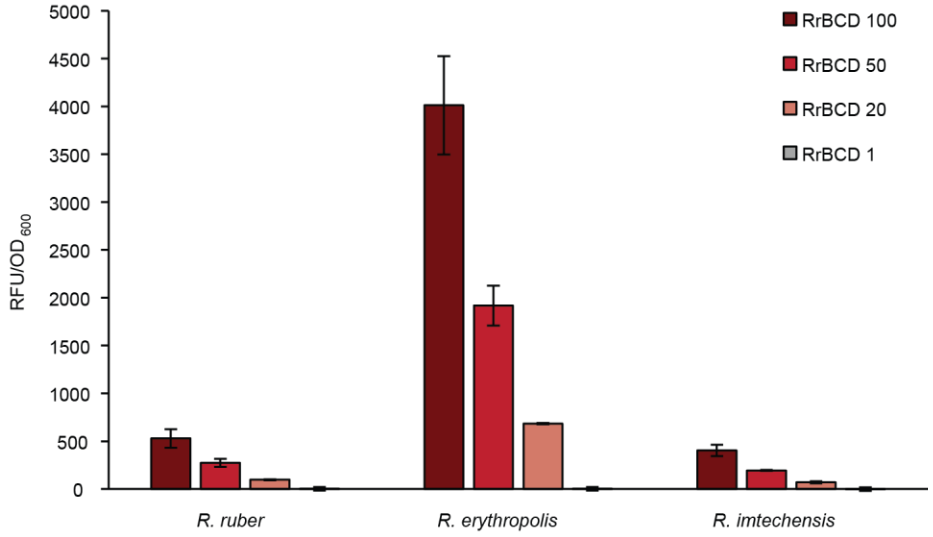


**Figure S7:** BCDs derived in *Rhodococcus ruber* maintain the expected rank order in additional *Rhodococcus* species. Data is plotted without normalization of absolute fluorescence across species.

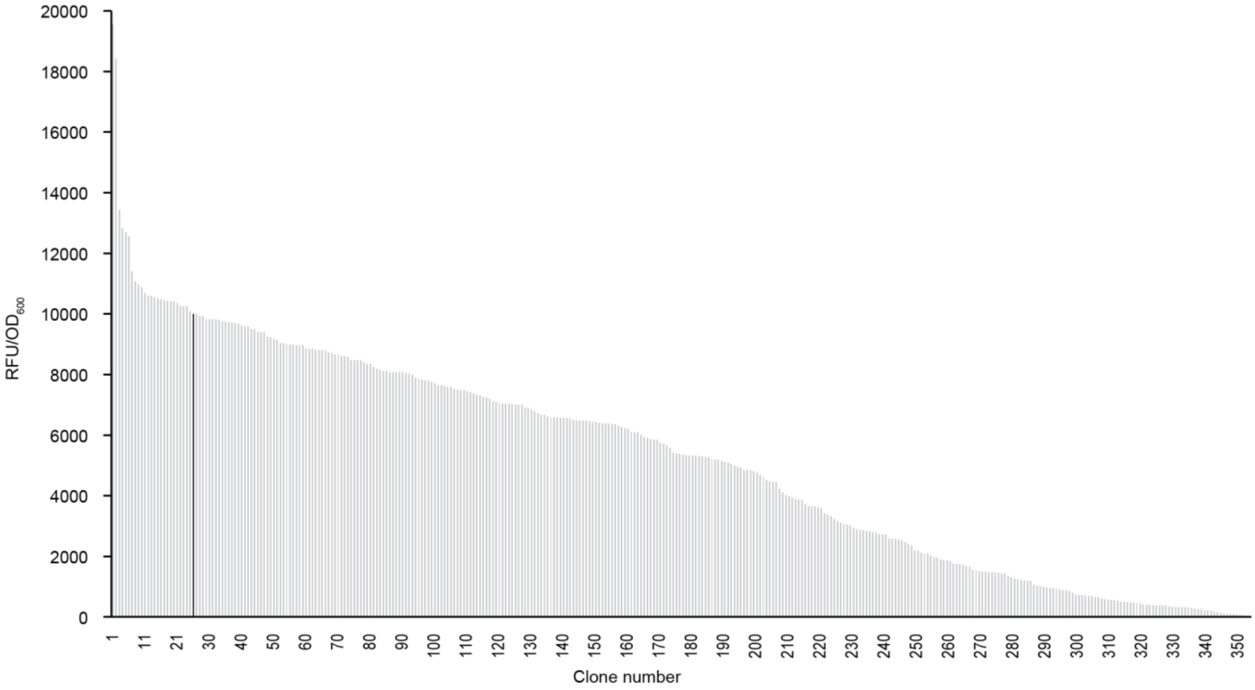


**Figure S8:** Distribution of 352 fully randomized (N9) SD2 variants screened using a weak promoter (P10). The strongest BCD, 25K, under the control of a weak promoter was included as a control (black).

**Supplementary Table 1**

The complete annotated sequence of the new BCD layout for *E. coli*. RiboJ is highlighted in magenta, the leader peptide 5´ UTR is highlighted in yellow with the SD1 sequence underlined and in bold, the leader peptide is highlighted in red with the SD2 region underlined and in bold, and the start codon is highlighted in cyan. The 4 nt cloning overhangs at the 5´ and 3´ ends are underlined and in bold. The 13 different 9 nt SD2 sequences corresponding to the different BCD translation initiation strengths are indicated.

| | |
|---|---|
| **GGAG**AGCTGTCACCGGATGTGCTTTCCGGTCTGATGAGTCCGTGAGGACGAAACAGCCTCTACAAATAATTTTGTTTAAGGGCCCAAGTTCACTTA**AAAAGGAGA**TCAACAATGAAAGCAATTTTCGTACTGAAACATCTTAATCATGC**NNNNNNNNN**TTAAA**TATG** | |
| SD2 25K: GGTAAGGAG | SD2 2.3K: TATGTGTTT |
| SD2 23K: GGAGGCAGC | SD2 1.5K: CGTCAAAAT |
| SD2 18K: TTGCAGAGG | SD2 0.75K: GGATTCTAG |
| SD2 13K: TTATCGGGG | SD2 0.45K: TCATGACCC |
| SD2 8.5K: GGCGCGGTG | SD2 0.2K: TCACGTCCC |
| SD2 4.6K: GCCGGTGTT | SD2 0.1K: TCAGGCCCC |
| SD2 3.75K: GGATTCTTG | |

**Supplementary Table 2**

The complete annotated sequence of the new BCD layout for *Rhodococcus*. RiboJ is highlighted in magenta, the leader peptide 5´ UTR is highlighted in yellow with the SD1 sequence underlined and in bold, the leader peptide is highlighted in red with the SD2 region underlined and in bold, and the start codon is highlighted in cyan. The 4 nt cloning overhangs at the 5´ and 3´ ends are underlined and in bold. The four different 9 nt SD2 sequences corresponding to the different BCD translation initiation strengths are indicated.

| | |
|---|---|
| **GGAG**AGCTGTCACCGGATGTGCTTTCCGGTCTGATGAGTCCGTGAGGACGAAACAGCCTCTACAAATAATTTTGTTTAAGGGCCCAAGTTCACTTA**ACTAAGGAA**TCAACAATGACCATGATTACGGATTCACTGGCCGTCGTTTTAGC**NNNNNNNNN**TTAAA**TATG** | |
| SD2 Rr100: AGAAGGAGA | SD2 Rr20: GCATTGCGG |
| SD2 Rr50: GGAGGCAGC | SD2 Rr1: TAGGACCCT |
